# Pitfalls in Inferring Human Microbial Dynamics from Temporal Metagenomics Data

**DOI:** 10.1101/073254

**Authors:** Hong-Tai Cao, Travis E. Gibson, Amir Bashan, Yang-Yu Liu

## Abstract

Human gut microbiota is a very complex and dynamic ecosystem that plays a crucial role in our health and well-being. Inferring microbial community structure and dynamics directly from time-resolved metagenomics data is key to understanding the community ecology and predicting its temporal behavior. Many methods have been proposed to perform the inference. Yet, we point out that there are several pitfalls along the way, from uninformative temporal measurements to the compositional nature of the relative abundance data, focusing on highly abundant species by ignoring or grouping low-abundance species, and implicit assumptions in various regularization methods. These issues have to be seriously considered in ecological modeling of human gut microbiota.

## I. INTRODUCTION

We coexist with trillions of microbes that live in and on our bodies [1]. Those microorganisms play key roles in human physiology and diseases [2]. Propelled by metagenomics and next-generation DNA sequencing technologies, many scientific advances have been made through the work of large-scale, consortium-driven metagenomic projects [3, 4]. Despite these technical advances that help us acquire more accurate organismal compositions and metabolic functions, little is known about the underlying ecological dynamics of our microbiota. Indeed, the microbes in our guts form very complex and dynamic ecosystems, which can be altered by diet change, medical interventions, and other factors [5–7]. The alterability of our microbiota not only offers a promising future for practical microbiome-based therapies [6, 8], such as fecal microbiota transplantation (FMT) [9, 10], but also raises long-term safety concerns. After all, due to its high complexity, careless interventions could shift our microbiota to an undesired state with unintended health consequences. Consequently, there is an urgent need to understand the underlying ecological dynamics of our microbiota; in the absence of this knowledge we lack a theoretical framework for microbiome-based therapies in general.

Inferring system dynamics is a typical task in system identification [11]. Measured temporal data, reasonable dynamical models, and objective criterion for model selection are the key elements in successfully inferring any system dynamics [12]. In the context of human gut microbiota, the measured temporal data are the time-series of microbe abundances, which are typically measured from the stool samples of a few individuals. Different dynamical models have been used to describe the dynamics of microbial ecosystems, e.g., linear models [13]; nonlinear models such as different variations of the Generalized Lotka-Volterra (GLV) model [14–19]; and other models [20]. Among theses models, GLV is a very popular one due to its simplicity. Given the measured temporal data and a dynamical model with many unknown parameters, we need to identify those parameters that yield the best model estimation according to certain criteria (e.g., minimum estimation error).

There are many existing methods to infer the microbial dynamics and reconstruct the ecological network from temporal metagenomics data based on the GLV model [21–24]. An overview of the workflow is depicted in Fig. 1. We apply certain perturbations to the systems (for example the administration of antibiotics or prebiotics) and measure the species abundances as a function of time using DNA sequencing technologies. What we don’t know is the underlying microbial dynamics, which can be parameterized in a population dynamics model with various model parameters, i.e., intrinsic growth rates, inter- and intraspecies interactions in the GLV model. In particular, the inter-species interactions can be captured by an ecological network and visualized as a directed graph shown in Fig. (1C). If the measured temporal data, i.e., the time-series of microbe abundances, are “rich” or informative enough, then we can reconstruct the ecological dynamics by identifying all the model parameters. The model parameters can then be used in turn to predict the temporal behavior of the microbial ecosystem, an ultimate goal of ecological modeling of human gut microbiota.

**FIG. 1.**
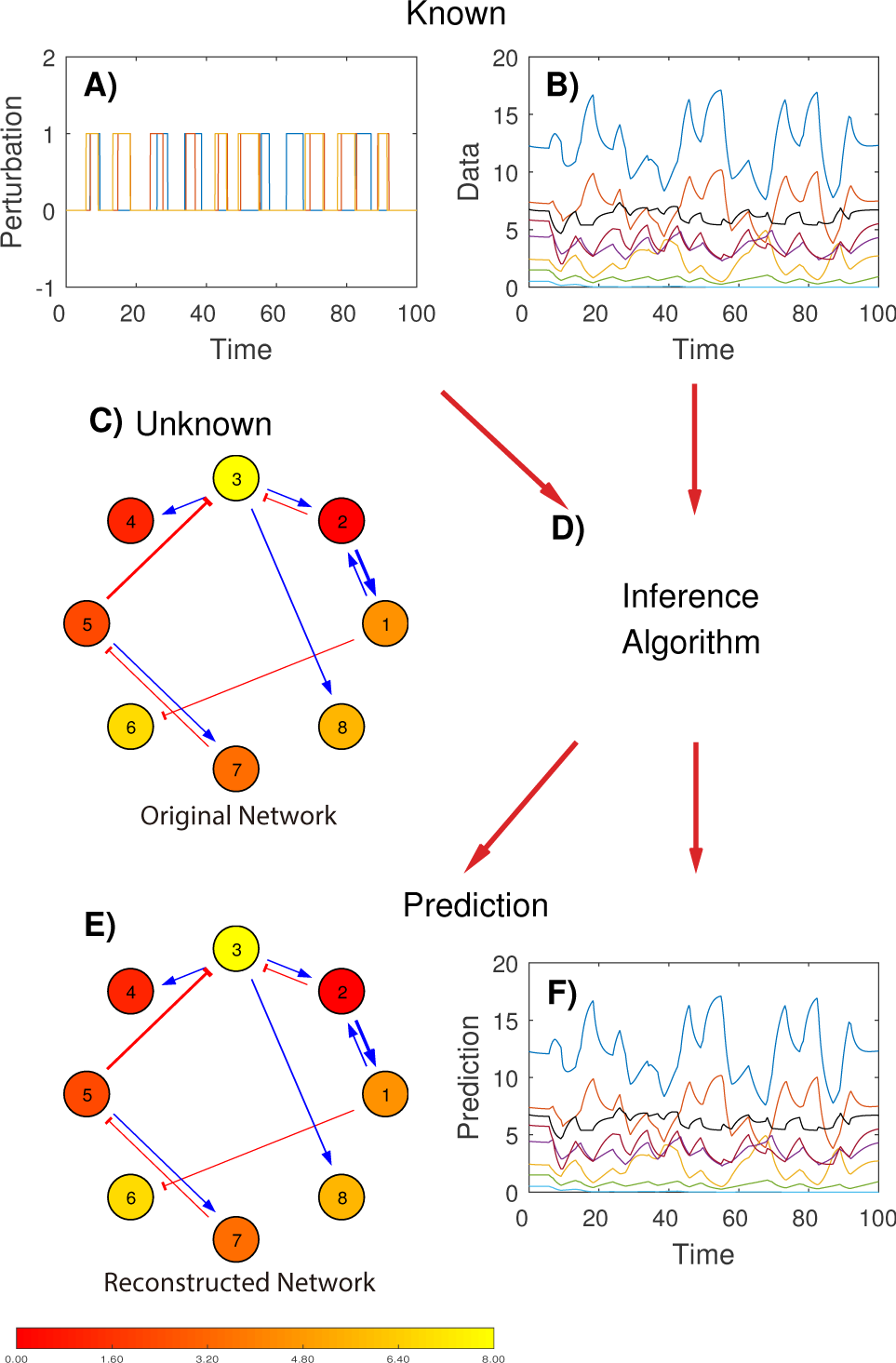
Overview of the workflow inferring microbial dynamics from time-series data. Given suitable perturbations (**A**) on a microbial ecosystem, and the corresponding time-series of microbe abundances (**B**), we aim to infer the microbial dynamics and reconstruct the underlying microbe-microbe interaction network (**C**), using classical population dynamics models, e.g., the Generalized Lotka-Volterra (GLV) model, and various standard system identification techniques (**D**). In the ideal case, the reconstructed microbe-microbe interaction network (**E**) captures all the key features of the original network (**C**), and the predicted time-series (**F**) agrees well with the original measurement (**B**). Yet, as pointed in this paper, there are many pitfalls in inferring the microbial dynamics from time-series data. In both (**C**) and (**E**), positive (or negative) interactions are shown in blue (or red) arrows, respectively. The absolute interaction strengths are proportional to the arrow widths and the microbiota growth rates are represented by circle colors.

Yet, this is just an ideal case. In reality, there are many pitfalls along the way. For example, the temporal data could be uninformative due to either low sampling rate or “unexcited” system dynamics. The compositionality nature of the relative abundance data will cause fundamental limitations in inference. And overlooking low-abundance but strongly interacting species might lead to erroneous model parameters. Those pitfalls are often ignored but can seriously affect the inference results. In this work we systematically study those pitfalls and point out possible solutions. Note that here we aim to reconstruct the ecological dynamics and the corresponding directed inter-species interaction network, rather than constructing any undirected microbial association network using similarity-based techniques, e.g., Pearson or Spearman correlations for abundance data or the hypergeometric distribution for presence absence data. The construction of microbial association networks has its own pitfalls, as discussed with detail in [25].

## II. KEY ELEMENTS IN INFERENCE

### A. Model

One of the key elements in system identification is choosing a reasonable dynamics model. Recently, population dynamics models, especially the classical GLV model, have been used for predictive modeling of the intestinal microbiota [17, 21–24]. Consider a collection of *n* microbes in a habitat with the population of microbe *i* at time *t* denoted as *x_i_*(*t*), the GLV model assumes that the microbe populations follow a set of ordinary differential equations (ODEs)

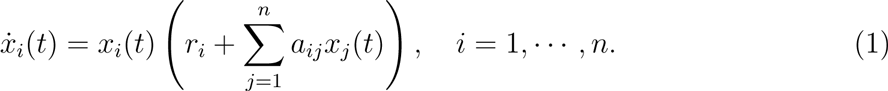

Here *r_i_* is the intrinsic growth rate of microbe *i*, *a_ij_* (when *i* ≠ *j*) accounts for the impact that microbe *j* has on the population change of microbe *i*, and the terms 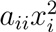 are adopted according to Verhulst’s logistic growth model [26]. Both *r_i_* and *a_ij_* are assumed to be time-invariant, i.e., they are constant regardless of how the system evolves over time. By collecting the individual populations *x_i_*(*t*) into a state vector 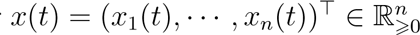, Eq. (1) can be represented in the compact form

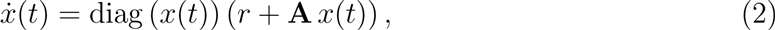

where *r* = (*r*_1_, …, *r_n_*)^┬^ ∈ ℝ^*n*^ is a column vector of the intrinsic growth rates, **A** = {*a_ij_*} ∈ ℝ^*n*×*n*^ is the inter-species interaction matrix, and diag generates a diagonal matrix from a vector.

The original GLV model, Eq. (2), excludes all the external perturbations applied to the system. For a class of asymptotically stable microbial ecosystems that follow this deterministic model and without any external perturbations, the microbe abundance profile will asymptotically approach a unique steady state [19]. The time-series data at steady state are rather uninformative and not very useful for system identification.

To excite the system and get “richer” or more informative time-series data, we have to apply external perturbations to drive the system and measure its response. In fact, we have to design very clever drive-response experiments to infer the underlying dynamics [20, 27]. Recently, an extended GLV model has been proposed to explicitly consider the impact of various external stimuli or perturbations *u_q_*(*t*)’s on the system dynamics [21, 23]:

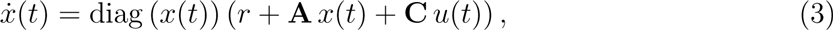

where *u*(*t*) = (*u*_1_(*t*), …, *u_l_*(*t*))^┬^ ∈ ℝ^*l*^ is the perturbation vector at time *t*, **C** = {*c_iq_*} ∈ ℝ^*n*×*l*^ is the susceptibility matrix with *c_iq_* representing the stimulus strength of perturbation *u_q_* on species *i*. This mimics realistic perturbations from antibiotics or prebiotics, which can inhibit or benefit the growth of certain microbes. The presence or absence of the antibiotics or prebiotics is evaluated as a binary perturbation *u*(*t*) (Fig. 1A) and the overall influences on the microbial species can be represented by the sum of products of susceptibility **C** and species abundance. We can then infer the microbial system under this particular drive-response scheme.

Besides the binary perturbation scheme, there is another type of drive-response experiment, which does not require us to introduce the susceptibility matrix C in the GLV model at all. This driving perturbation is implemented by setting up different initial conditions for the microbial ecosystem. For each initial condition change (which mimics the immediate result of an FMT), the system will respond by displaying certain transient behavior before it reaches the equilibrium (steady) state. If we concatenate several perturbed time-series corresponding to different initial conditions, we can treat the initial conditions as jumps or finite pulses from the equilibrium state. By construction, the concatenated time-series data contains various transient behavior of the system corresponding to different finite pulses, which could be very informative and help us infer the underlying system dynamics. Further comparisons between the above two drive-response experiments are discussed later (see Supplementary Figure 1).

### B. Data

Prior to the era of high-throughput DNA sequencing, microbiology studies heavily relied on cultivating microbes from collected samples. Yet, this process is rather tedious and time-consuming. Thanks to the development of next generation sequencing, we can now study microbiomes by direct DNA sequencing. In particular, the 16S ribosomal RNA (rRNA) gene targeted amplicon sequencing is a popular approach. In this approach, part of the 16S rRNA gene, which is the most ubiquitous and conserved marker gene of the bacterial genome, is sequenced [28]. Due to its simplicity, relatively low cost, and the availability of various developed analysis pipelines, this approach has become routine for determining the taxonomic composition and species diversity of microbial communities [29]. By filtering spurious reads and carefully clustering/grouping the remaining reads into the so-called Operational Taxonomic Units (OTUs) based on sequence similarity, one can obtain reliable and informative counts from 16S rRNA gene sequences. Indeed, as working names of groups of related bacteria, OTUs are intended to represent some degree of taxonomic relatedness. For example, when sequences are clustered at 97% sequence similarity, each resulting cluster is typically thought of as representing a biological species. One can then assign a frequency to each distinct genome within the microbial community describing their relative abundances within the population.

Note that comparing microbial composition between two or more populations on the basis of OTUs in their corresponding samples is totally different from comparing the absolute abundance of the taxa in the microbial ecosystems from which the samples are collected. Simply because we don’t know the total taxa abundance of the entire microbial ecosystem, it is only reasonable to draw inferences regarding the relative abundance of a taxon in the ecosystem using its relative abundance in the collected sample. In short, the microbial community can be described in terms of which OTUs are present and their relative abundances. The intrinsic compositionality of the relative abundance data will cause trouble in inference, as we will discuss in Sec. IIIC.

To reveal the pitfalls in inference, in this work we generate synthetic time-series data of microbe abundances using the classical GLV model. There are several advantages of using synthetic data. First of all, in this way we know exactly all the model parameters, and we can compare the inferred model parameters with their corresponding ground truth values. Second, we can freely choose the sampling rate and systematically study its impact on the performance of inference. Third, we can test different drive-response perturbation schemes and check which scheme offers more informative time-series data. Fourth, we can compare the difference of using absolute abundances and relative abundances in inference. Finally, we can compare the results of different regularization methods to check if certain regularization techniques perform better than others.

Since there is no closed-form solution to the ODEs of the GLV model (3), we solve them at predetermined time points. Many numerical integration methods such as explicit Runge-Kutta formula [30, 31], Adams-Bashforth-Moulton method [32] and Gear’s method [33, 34] can be used to approximate the solutions of (3). In this work, we choose the frequently used Runge-Kutta method. The total number of the synthetic data points is obtained by dividing the integral interval by the step-size. Note that the integral interval [0, *t*] in the numerical integration can be mapped to any length of time in reality, such as several weeks, days, or hours. To assign a realistic time unit to the synthetic data, we leverge two observations: (1) in our simulations (with the model parameters and initial conditions chosen in the way as described in Supporting Information), the GLV systems typically reach equilbrium state around t = 1; (2) after small perturbations human microbial ecosystems relax to the equilibrium state in about 10 days [17, 21, 24, 35]. Hence, we map the integral interval [0, *t*] in the simulation to [0, 10*t*] days in real time. For example, if we run the numerical integration from *t* = 0 to 10, this is equivalent to collecting the time-series data from day 0 to day 100. We emphasize that all the result presented in this work do not depend on the details of the time unit chosen in our simulations.

### C. Inference Methods

Let *x_i_*(*t_k_*) be the population of the *i*-th microbial species or OTU and *u_q_*(*t_k_*) be the *q*-th external perturbation at time point *t_k_*. Here, *k* = 0,1, …, *T*. The synthetic temporal data are generated based on the intrinsic growth rate vector r, the inter-species interaction matrix **A**, and the susceptibility matrix **C**. We need an *inference method* to identify all the model parameters in *r*, **A** and **C**, based on the time-series data {*x_i_*(*t*),*u_q_*(*t*)}.

Move *x_i_*(*t*) of (3) to the left hand side and then integrate both sides over the time interval [*t_k_*,*t*_*k*+1_), yielding

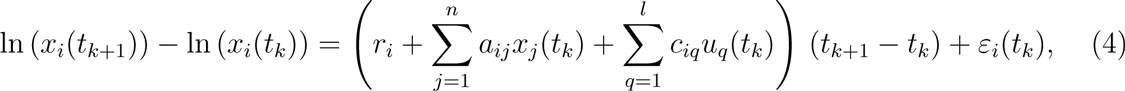

where we have assumed that *x_i_*(*t*) and *u_p_*(*t*) are roughly constant over *t* ∈ [*t_k_*, *t*_*k*+1_), *t_k_* ≥ 0. Here *ε_i_*(*t_k_*) represents the corresponding error arising from the approximation of the integral by holding the integrand constant over the time interval.

Define the scaled log-difference matrix **Y** = {*y_ik_*} = {*y_i_*(*t_k_*)} ∈ ℝ^*n*×*T*^ where *y_i_*(*t_k_*) = [ln(*x_i_*(*t*_*k*+1_))−ln(*x_i_*(*t_k_*))]/(*t*_*k*+1_−*t_k_*), the parameter vector 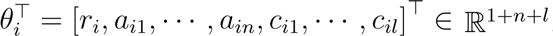, and the vector *ϕ_k_* = [1,*x*_1_(*t_k_*), …, *x_n_*(*t_k_*), *u_l_*(*t_k_*), …, *u_l_*(*t_k_*)]^┬^ ∈ ℝ^1+*n*+*l*^, then the discretized GLV model (4) can be represented by a system of linear algebraic equations:

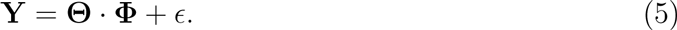

Here 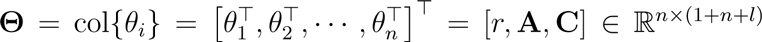 is the parameter matrix that needs to be identified. *∊* ∈ ℝ^*n*×*T*^ represents the corresponding approximation error matrix. Φ = row{*ϕ_k_*} = [*ϕ*_0_,*ϕ*_1_, …,*ϕ*_*T*−1_] ∈ ℝ^(1+*n*+*l*)×*T*^. Equation (5) is often called the *identification function* that can be used to solve for the unknown parameter matrix Θ.

Given any time-series data *x*(*t_k_*) and *u*(*t_k_*) of the GLV model, Θ should be a solution of the identification function (5). Yet, Θ usually cannot be exactly solved, since (5) is usually *underdetermined* because of the limited available data. Indeed, the number of equations *n* × *T* is typically less than the number of unknowns *n* × (1 + *n* + *1*). Θ can be approximately solved by optimization methods. There are many algorithms to obtain an approximate solution, though. We discuss those methods as follows.

#### 1. Least square

Mathematically, Θ can be estimated as 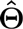 by solving the following optimization problem:

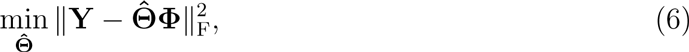

where 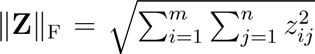 is the Frobenius norm of matrix **Z** = {*z_ij_*} ∈ ℝ^*m*×*n*^. The solution 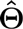 can be obtained by the classical least-square regression method:

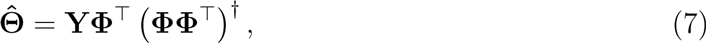

where (ΦΦ^┬^)^†^ represents the pseudo-inverse matrix of ΦΦ^┬^. Note that (ΦΦ^┬^)^†^ = (ΦΦ^┬^)^−1^ when ΦΦ^┬^ is non-singular.

#### 2. Regularizations

In statistic regressions, the least-square solution (without any penalty) (7) can be biased and cause overfitting, which results in extremely large absolute values in the model parameters, rendering the results meaningless. It is possible that the matrix ΦΦ^┬^ is nearly singular, which will cause its inverse matrix numerically unstable and the estimation over-fitted. To avoid extreme parameter values and increase the numerical stability of the inference algorithm, we can add penalty terms to the regression. This is often called *regularization* in regression methods.

Depending on the penalty terms (e.g., based on *ℓ*^1^-or *ℓ*^2^-norm), there are many di.erent types of regularization methods. In particular, lasso regularization [36.38], which uses *ℓ*^1^-norm penalties, solves the regression problem in the form of

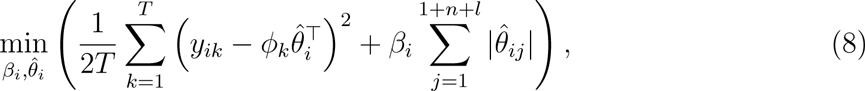

where 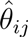 is the *j*-th element in 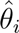 and *i =* 1,2, …,*n*. Lasso regression estimates the unknown parameters in the *i*-th row of 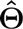. There are several algorithms solving this optimization problem, such as truncated singular value decomposition, l-curve, cross validation and so on. Detailed algorithms and discussions can be found in [39]. In this work, we use the *k*-fold cross validation method and let *k* = 5 in lasso regularization.

To perform cross validation, we divide the the entire data into two parts: *training* and *test.* The training data is used to solve the optimization problem and identify the model parameters. The test data is used for validation. In *k*-fold cross validation, we divide the original data into *k* data sets, randomly choose one set as the test data, and use the rest (*k* − 1) data sets as the training data. The regularizations are performed on the training dataset, and then the residual error of the test set (or the whole dataset) is constructed as a function of *β_i_*. This function is used to measure the accuracy of time-series prediction, where smaller error means better estimation. The value of *β_i_* is set when this function reaches its minimum. Finally the estimation of the model is obtained by the regularization over the total dataset.

Different from the lasso regularization that uses *ℓ*^1^-norm penalties, Tikhonov regularization, as known as ridge regression in statistics, uses *ℓ*^2^-norm penalties:

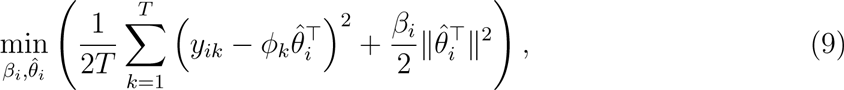

where ||·|| represents the *ℓ*^2^-norm and *i* = 1,2, …, *n*. Similar to lasso regression, the above penalty terms *β_i_* can also be determined by cross validation. There are *n* different *β_i_*’s penalizing all the model parameters. Alternatively, we can let the same *β_i_* penalize the same part of the estimations 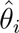 in 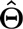 so that the growth rate vector *r*, interaction matrix **A** and susceptibility matrix **C** are penalized separately [21]:

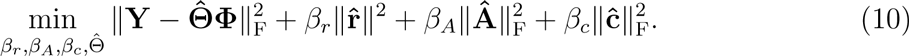

Note that in the original optimization problem (9), it is not required that some *β_i_*’s should be the same value mathematically. For simplicity, the estimation 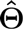 is penalized by using different *β_i_* in two different schemes: vector scheme and matrix scheme. The former is an optimization problem in the form of (9) by breaking up the matrix 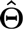 into vectors and using different penalty terms for different rows of 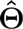. The latter treats the *r*, **A** and **C** as separate objects and use the same *β_i_* for penalizing the same object. It then solves the regularization in the form of (10), as shown above. As there is an implicit assumption that the *r*, **A** and **C** should be penalized separately in (10), it adds more constraints when identifying the parameters in the extended GLV model. We prefer the original optimization (9) because of its flexibility in choosing the penalty terms.

Linear combinations of *ℓ*^1^- and *ℓ*^2^-norm penalties in (8) and (9) result in the so-called elastic net regularization method [40]:

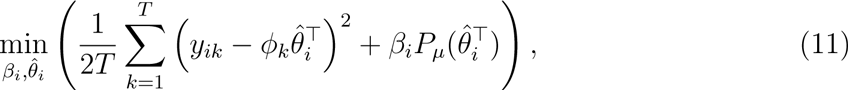

where 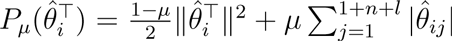, and *μ* ∈ [0, 1] is a predetermined parameter for the optimization. The elastic net regularization becomes the Tikhonov (or lasso) regularization when *μ* = 0 (or 1), respectively.

All the regularization methods (lasso, Tikhonov and elastic net) use penalty terms to regularize the least-square regression. The penalty terms make the absolute values of estimation smaller and suppress the unimportant parameters to 0. Due to the presence of penalty terms 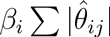 in lasso regularization (8), unimportant parameters in *θ_i_* will be forced to be 0. Therefore lasso is a kind of *sparse* regression that implicitly assumes the interaction matrix **A** in the GLV model is sparse (which is of course not necessarily true). By contrast, Tikhonov regularization (9) does not force the parameters to be zero, because the penalty term 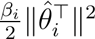 does not penalize the absolute values in the estimation. Therefore Tikhonov regularization does not have the property of sparse regression. Although these regularization methods reduce the norm of estimation and aim to make the results more realistic, it doesn’t mean the results are getting close to the ground truth, as we will discuss in Sec. IIIE.

## III. PITFALLS

### A. Accurate time-series prediction does not imply accurate inference

Since the ground truth is typically unknown in real-world system identification problems, the identified system parameters are usually verified by simulating the model dynamics and comparing the predicted time-series with the measured one. This is suitable for simple systems with the number of unknown parameters much smaller than the number of data points. Yet, for complex ecosystems such as human gut microbiota that have hundreds of species, this approach is rather unreliable in practice. Consider the popular binary perturbation scheme of microbial systems described by the extended GLV model (3), which is a system of nonlinear differential equations with *n*(1 + *n* + *l*) unknown parameters for *n* species with *l* external perturbations. Since the time-series data typically have very limited time resolution and very few data points, we are facing an underdetermined problem (the number of equations, which is proportional to the number of data points, is much less than the number of unknowns). Over-fitting is a notorious issue in this scenario. Even if the temporal behavior of microbial systems are predicted with high accuracy, there is no guarantee that the identified model parameters are close to their ground truth values. Indeed, accurate temporal predictions are possible even if the identified interactions look totally different from the actual ones [41].

To demonstrate the above point, we set up a synthetic microbial system with 8 species, following the GLV dynamics with 3 binary perturbations. It is a microbial system with homogeneous interaction strengths among all species with mean degree 6.4 in the underlying ecological network. The abundance of a certain species is increased when its susceptibility is positive and the binary perturbation is turned on. The population of all the species in the microbial systems are simulated from *t* = 0 to 10, mapped to 100 days. The sampling rate is set to be once per day, which means there are total 100 data points for this data set, where the time interval between two adjacent data points is one day.

Comparing Fig. 2A2 and A3, we find that we can accurately predict the temporal behavior of microbial population, given the same initial conditions and the time-series perturbation data (Fig. 2A1). Yet, the identified inter-species interaction network (Fig. 2B2) looks drastically different from the ground truth (Fig. 2B1). For example, some strong interactions (e.g., 2 → 1) are lost, and some unessential interactions are inferred as dominant interactions (e.g., 6 → 5). In fact, all the identified model parameters are quite different from the ground truth (see Fig. 2C1-C3).

**FIG. 2.**
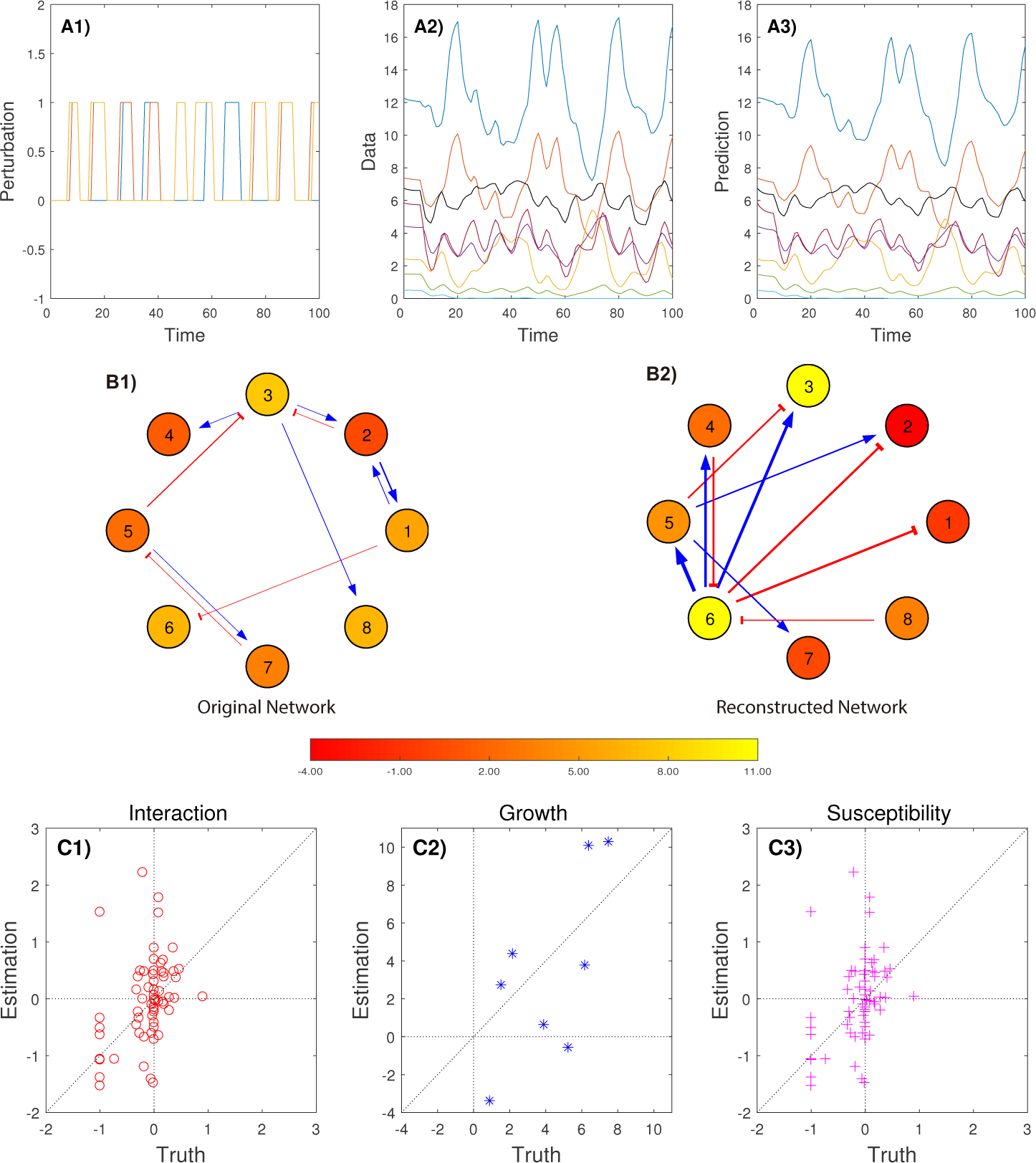
Perfect time-series prediction does not imply accurate network reconstruction. **A1**: Time series of binary perturbations. **A2**: Synthetic time-series of species abundances generated from a GLV model. Both perturbation and abundance data are sampled once per day. **A3**: Predicted time-series of species abundances calculated from the inferred GLV model. **B1**: Original inter-species interaction network. **B2**: Reconstructed inter-species interaction network. Here in both B1 and B2 only the top-10 strongest interactions are shown. Circle colors represent growth rates. **C1**: Inferred interaction strengths v.s. true interaction strengths. **C2**: Inferred growth rates v.s. true growth rates. **C3**: Inferred susceptibilities *v.s.* true susceptibilities.

The inference method used in generating Fig. 2 is just the least-square regression, which is fast and effective in predicting the population behavior. But it has a potential drawback of over-fitting, especially when there are a lot of estimation parameters while the given time-series is not long enough. Here, we have 100 data points and 96 model parameters, hence it is not a big surprise that there is an over-fitting issue and the regression results are not reliable. If the sampling rate is increased to 100 times per day (which is of course more challenging and expensive), then both the prediction of microbial populations and interaction network reconstruction become more accurate and the over-fitting issue will be largely mitigated, as shown in Fig. 1.

The above result clearly demonstrates that accurate temporal prediction could be just due to over-fitting, and the identified model parameters could be far from the ground truth.

### B. Sampling rate really matters

Different sampling rates capture different resolutions of the dynamics of the microbial system [42]. Inferred microbial networks from time series data can be misleading if the microbial system is sampled at an improper frequency. Unfortunately, there is no simple rules like nyquist frequency for the GLV model, and the ideal sampling rate depends on the particular microbial system of interest [42, 43]. Results presented in Fig. 1 and Fig. 2 clearly suggest that sampling rate is really an important factor determining the performance of inference, as discussed in detail below.

We are interested in the continuous-time dynamics of microbial communities. But measurements are always taken at discrete time points, resulting in discrete time-series data. The sampling rate becomes crucial as it bridges the measured discrete time-series data and the original microbial system. Obviously, higher sampling rate makes the interpolated discrete time-series data better approximate the continuous-time dynamics of the original system. It should be pointed out that the inference method (4) itself requires high sampling rates. The scaled log difference *y_ik_* represents the linearized approximation of the GLV, whose numerator is a nonlinear function and the denominator is the step size. The approximation error is *ε_i_*(*t_k_*) = *y_ik_* − *c*, where 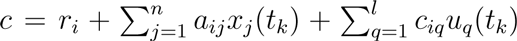 in the integral interval [*t_k_*,*t*_*k*+1_). As *t*_*k*+1_ − *t_k_* increases linearly, *y_ik_* changes nonlinearly, which results in a nonlinear increase of *ε_i_*(*t_k_*). Sampling rate becomes substantial because of this nonlinear behavior of the approximation error. Though we can arbitrarily increase the sampling rate for synthetic data, it is rather costly in real data collection. Hence, it would be more desirable if the time-series data can approximate the original microbial dynamics with higher accuracy at a lower sampling rate.

The binary perturbation scheme helps us excite the system to get more informative time-series data, but the extended GLV model (3) introduces more model parameters (which consist of the whole susceptibility matrix C) than the original GLV model (2). Moreover, the presence and absence of the introduced antibiotics or prebiotics bring new difficulties. More unknown model parameters apparently require time-series data with higher sampling rate. In reality, due to many limitations, the finest longitudinal data of human gut microbiota is actually sampled on just a daily basis. Hence, using the original GLV model with concatenated time-series with different initial conditions could be a better way to perform inference. Indeed, we find that this initial-condition-perturbation scheme is much better than the binary perturbation scheme in terms of smaller number of unknowns. It also provides higher accurate inferring results comparing to the binary external perturbations (see Supplementary Figure 1).

We now evaluate the impacts of sampling rate on the performance of inference with the initial-condition-perturbation scheme. We choose four sampling rates: weekly, every two days, daily, and twice a day, as shown in Fig. 3. Indeed, we find that the higher sampling rate, the better inference results. Even the data are sampled every two days, the inferred interactions are much more reliable than the results with weekly sampling rate.

**FIG. 3.**
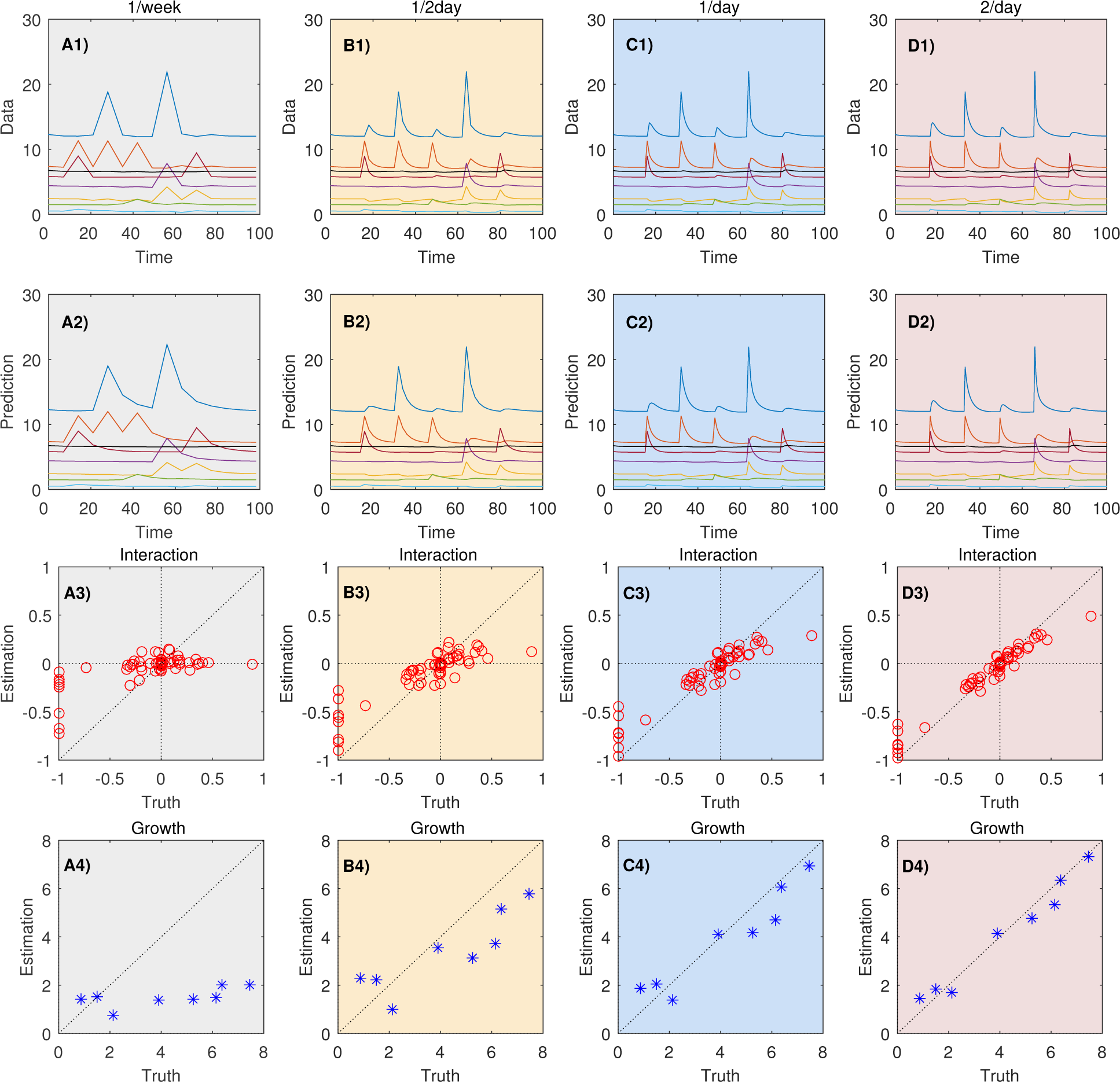
Impact of sampling rates on inferring microbial dynamics. Row-1: Time-series of species abundances generated from a GLV model with different sampling rates: (**A1**) once a week; (**B1**) every two days; (**C1**) daily; and (**D1**) twice a day. Row-2: Predicted time-series of species abundances calculated from the corresponding inferred GLV model. Row-3: True interaction strengths v.s. inferred interaction strengths from time-series data of different sampling rates. Row-4: True growth rates v.s inferred growth rates from time-series data of different sampling rates.

In reality, the initial-condition-perturbation can be implemented by fecal microbiota transplantation, which immediately changes the abundances of multiple species (or even introduces some new species). In the rest of this paper, we will focus on this type of perturbation.

### C. Compositionality raises serious challenges

As discussed in Sec.IIB, microbial communities can be typically described in terms of memberships and relative abundances of OTUs. Of course, if the total population is roughly time-invariant, then the compositionality of relative abundance data will not significantly alter the original time-series data. This may lead us to conclude that this will not affect the inference. Yet, this is not the case. A time-invariant total population will be linearly correlated with the constant row in Φ, introducing linear correlations of rows of Φ. This will leads to the rank deficiency of the matrix ΦΦ^┬^. Hence a roughly time-invariant total population will cause ΦΦ^┬^ to be almost singular, drastically reducing the numerical stability of the inverse and worsening the inference results.

It seems that there is no numerical stability issue if the total population is not time-invariant. However, this is not the case. To demonstrate the effect of compositionality of relative abundance data in inference, we normalize the original time-series data obtained from the same microbial system used in Fig. 3. We simply apply the least-square regression method to both original and normalized time-series data. Results are shown in Fig. 4. It is obvious that normalizing the original time-series data makes the inference results worse. Moreover, the accurate prediction of the microbial populations becomes impossible. These results can be explained by the fact that normalization makes ΦΦ^┬^ singular. Let 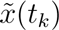 be the normalized data of *x*(*t_k_*). It holds that 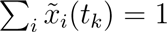. Denote 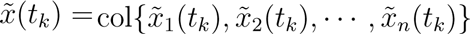, *k* = 0,1, … *T*–1. Let 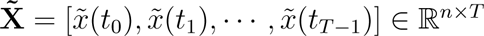. Apparently, the rows of 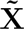 are linearly dependent, and 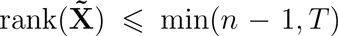. Let **1** ∈ ℝ^1×*T*^ be a row vector and **U** = [*u*(*t*_0_),*u*(*t*_1_), …,*u*(*t*_*T*−1_)] ∈ ℝ^1×*T*^, then 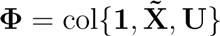. We have rank(Φ) ≤ min(*n* − 1 + *l,T*), simply because the constant row (i.e., the first row) and one row in the 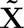 partition of Φ are linearly correlated with the other rows in Φ. This leads to rank(ΦΦ^┬^) ≤ *n* − 1 + *l* and thus ΦΦ^┬^ is rank deficient (i.e., singular).

**FIG. 4.**
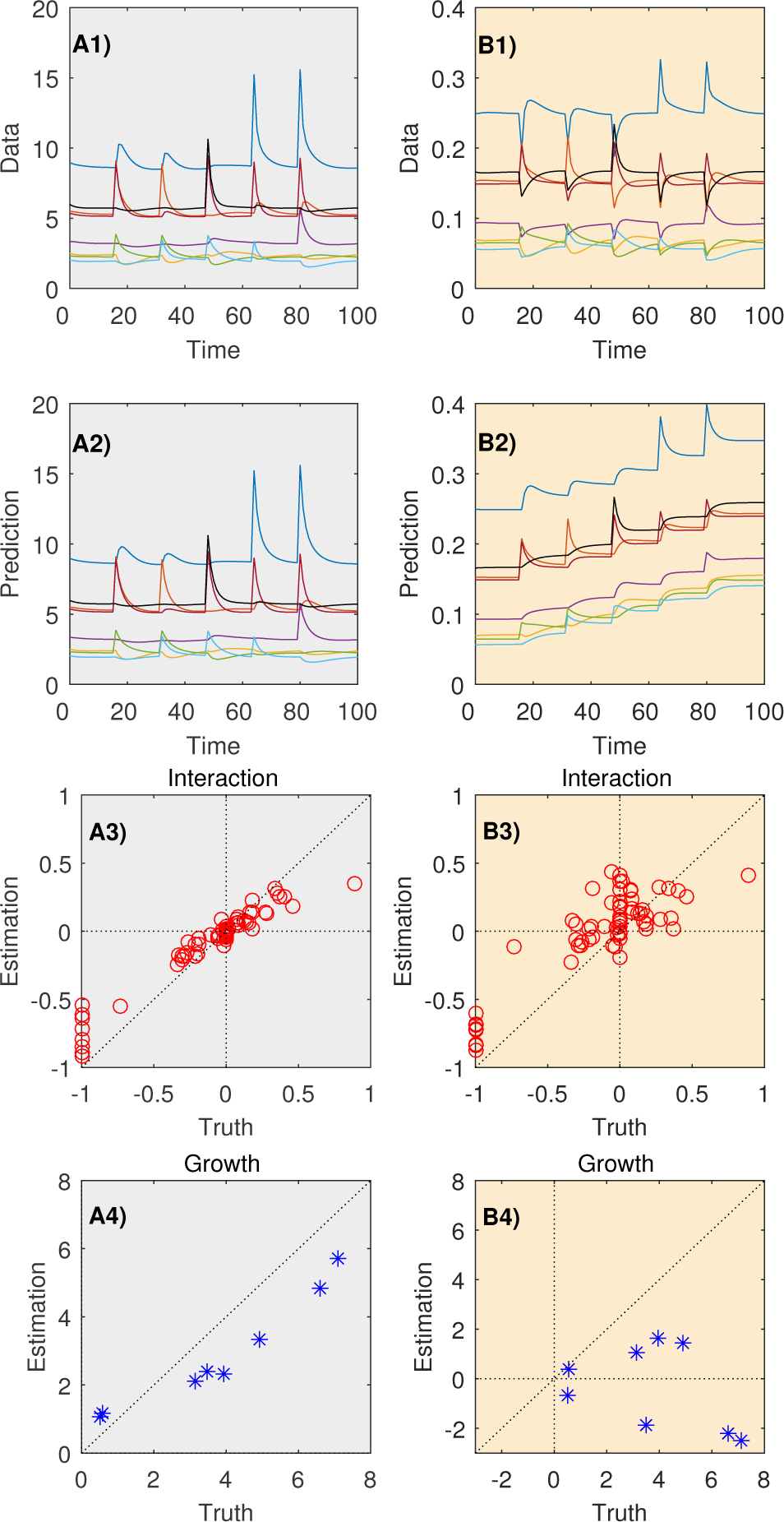
Compositionality of relative abundance data impedes the inference of microbial dynamics. Column-1: using absolute abundances data. (**A1**) Time-series of absolute abundances; (**A2**) Predicted time-series of absolute abundances; (**A3**) True interaction strengths v.s. inferred interaction strengths; (**A4**) True growth rates v.s. inferred growth rates. Column-2: using relative abundances data. (**B1**) Time-series of relative abundances; (**B2**) Predicted time-series of relative abundances; (**B3**) True interaction strengths v.s. inferred interaction strengths; (**B4**) True growth rates v.s. inferred growth rates. Inference results from relative abundances are far from the ground truth. The time-series prediction of relative abundances also differ significantly from that of the original relative abundances.

In addition to causing rank deficiency, compositionality will cause a more serious issue: distorting the original dynamics. As shown by the top (blue) curves in Fig. 4A1 and B1. The first jump is a positive jump in the original data (A1), representing an increase in absolute abundance of this species. Yet, it becomes negative after normalization (B1), indicating a decrease in the relative abundance of this species. This distortion of the oringinal temporal data will definitely affect the inference results. One promising solution to resolve this issue is to measure overall microbial biomass over time in the ecosystem via the quantitative PCR technique [21, 23, 24].

### D. Grouping or ignoring low-abundance species lacks justification

Since the number of equations is typically much smaller than the number of unknowns, many previous works group those low-abundance species together and treat them as a pseudo-species [21, 22]. This approach sounds very rational in reducing the number of unknowns (i.e., model parameters). Yet, we don’t know if it indeed works as we expected. In case the low-abundance species are also strongly interacting species (i.e. they interact strongly with their interacting partners), they can easily drive the the microbial ecosystem to different steady states [19]. Simply grouping all the low-abundance species together might generate distorted interaction networks. To test this idea, we systematically study the impact of grouping low-abundance species in inferences.

We define high-abundance species to be those species that account up to 90% of the total abundance or more in the sampled time series data. We compare three different scenarios: (1) We infer the interactions using the entire time series data without grouping low-abundance species. (2) We simply remove the low-abundance species in the temporal data, and focus only on the remaining species. (3) We group all the low-abundance species as a new species, and then perform the inference. Inspired by [19], we deliberately generate a microbial system with interaction strength heterogeneity. The inferred results for the above three scenarios are shown in Fig. 5. Note that when all the species are considered, the identified interactions are accurate. Yet, ignoring or grouping low-abundance species leads to poor inference results.

**FIG. 5.**
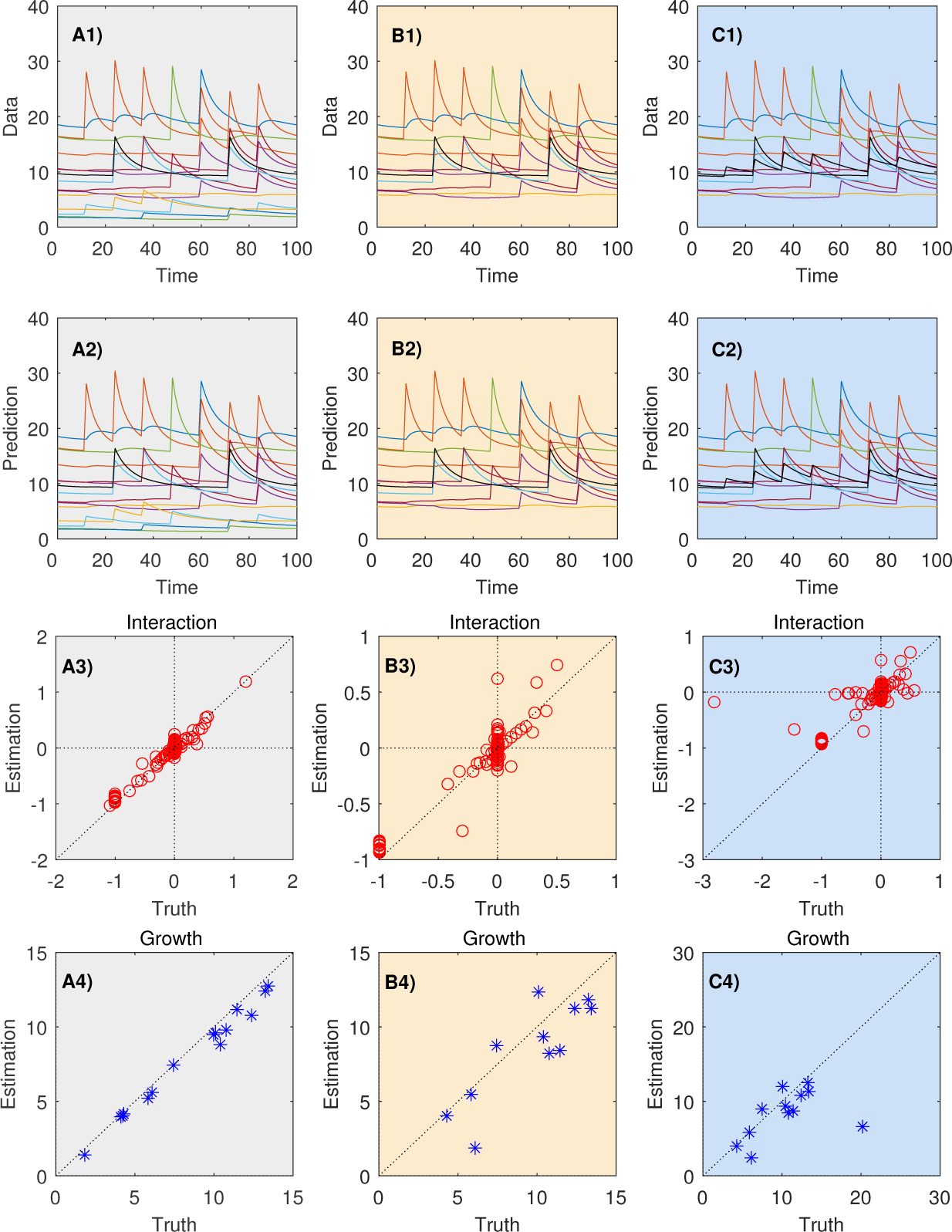
Ignoring or grouping low-abundance species impedes the inference of microbial dynamics. Column (**A**): Without ignoring or grouping of low-abundance species, the inference results are acceptable, and the predicted time-series agrees well with the original time-series data, provided the sampling rate is high enough. Column (**B**): After ignoring the low-abundance species, the inference results are much worse, despite the predicted time-series still agrees well with the original time-series data. Column (**C**): If we group the low-abundance species together and regard them as a new species, the inference results are still not comparable to the results of using original data. In generating these figures, we consider a system of *n* = 15 species with an heterogeneous inter-species interaction network with mean degree 〈*k*〉 = 11.2.

We emphasize that grouping low-abundance species is not a solution to the underdetermined problem. Even the microbial interaction network is assumed to be homogeneous, reconstructed network obtained by grouping some low-abundance species can be misleading, because grouping can create false interactions between the grouped species and highly abundant species.

### E. Regularizations need to be done with care

Since the identification function (5) is typically under-determined, to avoid over-fitting, regularization methods such as (9), (8) and (11) are preferred to the least-square regression method (no regularization) (7). To determine which of the methods: least-square regression (no regularization), Tikhonov (with *ℓ*^2^-norm penalty), lasso (with *ℓ*^1^-norm penalty) and elastic net (a linear combination of *ℓ*^1^- and *ℓ*^2^-norm penalties), works best, we apply them to the same time-series data (Fig. 6). We find that least-square regression certainly cannot identify the model parameters. To our surprise, Tikhonov regularization does not work well either. This is partially due to the fact that it penalizes the square of unknowns, rather than the absolute values of the unknowns as lasso regularization does. If the unknowns have orders of magnitude differences, then Tikhonov regularization is doomed to failure. By contrast, lasso regularization shrinks the absolute values of the unknowns to avoid the over fitting problem. Hence it works very well even if the unknowns could have orders of magnitude differences. Finally, although elastic net regularization combines both *ℓ*^1^- and *ℓ*^2^-norm penalties, there is no clear improvement in its inferred results.

**FIG. 6.**
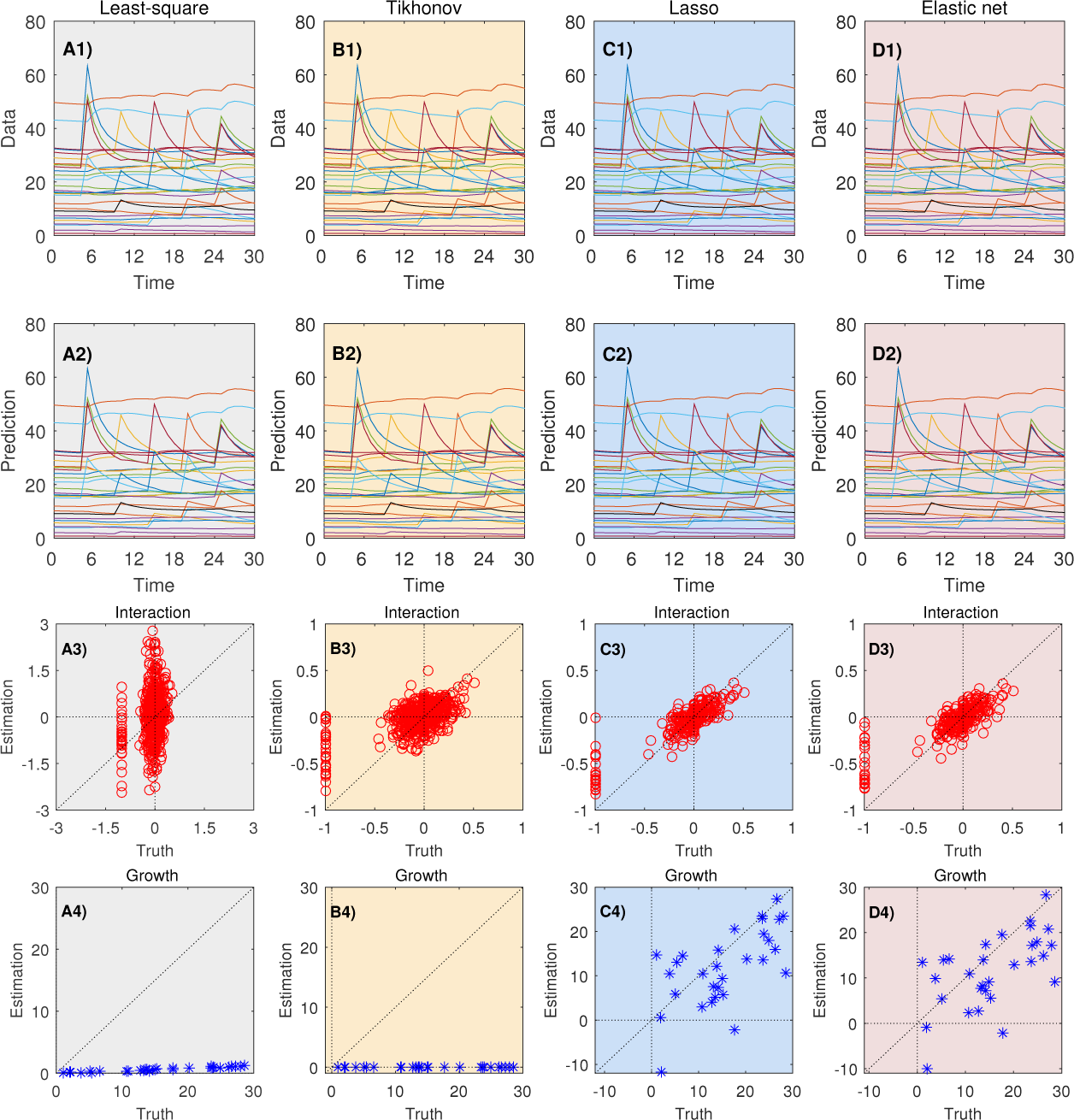
Inappropriate regularization impedes the inference of microbial dynamics. Column (**A**): Without any regularization, we can perform the inference using the least-square method (i.e., no penalty terms). The inference results are not acceptable. Column (**B**): With Tikhonov regularization (also known as *L*_2_-regularization or ridge regression), the inference results are still bad. Column (**C**): With lasso regularization (also known as *L*_1_-regularization), the inference results are slightly better. Column (**D**): With elastic net regularization, which uses a linear combination of *ℓ*^1^- and *ℓ*^2^-norm penalty terms (with *μ* = 0.5 in (11)), the inference results are as good as that of using lasso only. Note that in all the four cases, the predicted time-series agrees well with the original time-series data. In generating these figures, we consider a microbial ecosystem of *n* = 30 species with an homogeneous inter-species interaction network and mean degree 〈*k*〉 = 23.2.

## IV. CONCLUSIONS AND PROSPECTS

Inferring microbial dynamics from temporal metagenomics data is a very challenging task. Existing methods work well in predicting the population evolution of microbial systems. Yet, the identified model parameters might be totally different from their ground-truth values. Without direct experimental validation, it is hard to conclude that the inferred dynamics represents the true underlying microbial dynamics. New inference methods that can leverage some prior knowledge of the growth rates or/and inter-species interactions need to be developed.

Note that in this work, we do not focus on some other issues in dealing with real micro-biome data, e.g., measurement noise, which of course will also affect the inference. Instead, we focus on synthetic data generated from GLV model. We point out that even with “clean” time series data, current technological limitations and common practices can lead to poor system identification. Some of these pitfalls can be overcome with more information, i.e., the measurement of total bacterial biomass present in the samples using qPCR techniques. Other pitfalls are more difficult to deal with. New inference methods that can take full advantage of existing microbiome datasets still need to be developed. In particular, Bayesian inference algorithms could be very useful in practice, because they can not only estimate error in inferences of dynamical systems parameters but also perform statistical modeling of temporal metagenomics data [24].

## Acknowledgements

This work was partially supported by the John Templeton Foundation (award number 51977).

## Author Contributions

Y.-Y.L. conceived and designed the project. H.-T.C. performed all the numerical simulations and data analysis. All authors analysed the results. Y.-Y.L. and H.-T.C. wrote the manuscript. All authors edited the manuscript.

## Author Information

The authors declare no competing financial interests. Correspondence and requests for materials should be addressed to Y.-Y.L.

## SUPPORTING INFORMATION

### A. Generate the synthetic data

We use a class of asymptotically stable GLV models for the generation of synthetic data [1]. Let **N** be a random matrix whose elements are drawn from the standard normal distribution, i.e., 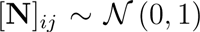. Let **H** be a diagonal matrix whose diagonal elements are either all ones or from a power-law distribution, i.e., 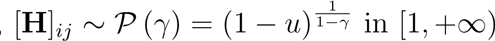 with exponent *γ* = 1.2, and *u* taken from a uniform distribution 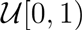. Let 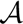 be the adjacency matrix of the interaction network in the microbial community. Finally, the interaction matrix **A** is generated by

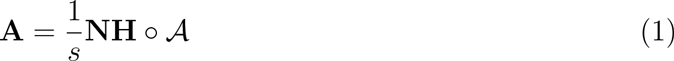

with all diagonal elements *a_ii_*’s set to be −1 to ensure stability. Here, *s* is a scaling factor, and the Hadamard product o multiplies two matrices of the same dimension element by element. If the diagonal matrix **H** has all ones for diagonal elements, then the columns of **A** have similar mean absolute values and all the species are equally interactive, which generates a homogeneous interaction network. If the diagonal elements of **H** are taken from a power-law distribution, then some columns will have high mean absolute values and the corresponding species become the so-called strongly interactive species, which generates a heterogeneous interaction network. 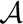 determines the total number of interactions among all the species in the microbial community, whose off-diagonal elements follow a Bernoulli distribution, i.e., 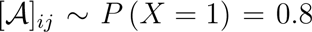. To ensure stability, the scaling factor *s* = 1 if the **H** generates a homogeneous interaction network, otherwise 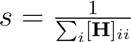 [1].

Regarding the intrinsic growth rate vector *r*, if we allow for low-abundance species, we draw [*r*]_*i*_ from a uniform distribution 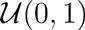. Otherwise, we simply choose 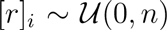, where *n* is the total number of species. In both cases, we keep generating intrinsic growth rate *r* until the expected steady state *x*^*^ = −**A**^−1^*r* becomes feasible, i.e., 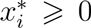 for all species. For the susceptibility matrix **C**, we choose 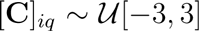. In the case of binary perturbation scheme, we choose the initial conditions at *t*_0_ to be close to the equilibrium state, i.e., [−**A**^−1^*r*, − 1.05**A**^−1^*r*], so that the intrinsic system relaxation doesn’t mix up with its response to external binary perturbation. In the case of changing initial conditions scheme, we use the same initial conditions as the case of binary perturbation scheme. Finally the GLV model is solved according the Runge-Kutta method. The final time-series data are sampled as needed from the numerical integration results.

### B. Compare the drive response model

To compare the inference results of two driving-response schemes: (1) external binary perturbations; (2) changing of initial conditions, we apply them to synthetic data generated from the same microbial system. We apply the least-square regression method to infer the dynamics at two different sampling rates. Details are shown in Supplementary Fig. 1. We find that concatenating time-series of different initial conditions yields much better inference results than external perturbations. Indeed, the identification results of the external binary perturbation scheme look really bad. Yet, its predicted time-series still agrees well with the original one, partially because the susceptibility matrix C is inferred properly in this case.

---

**FIG. 1.**
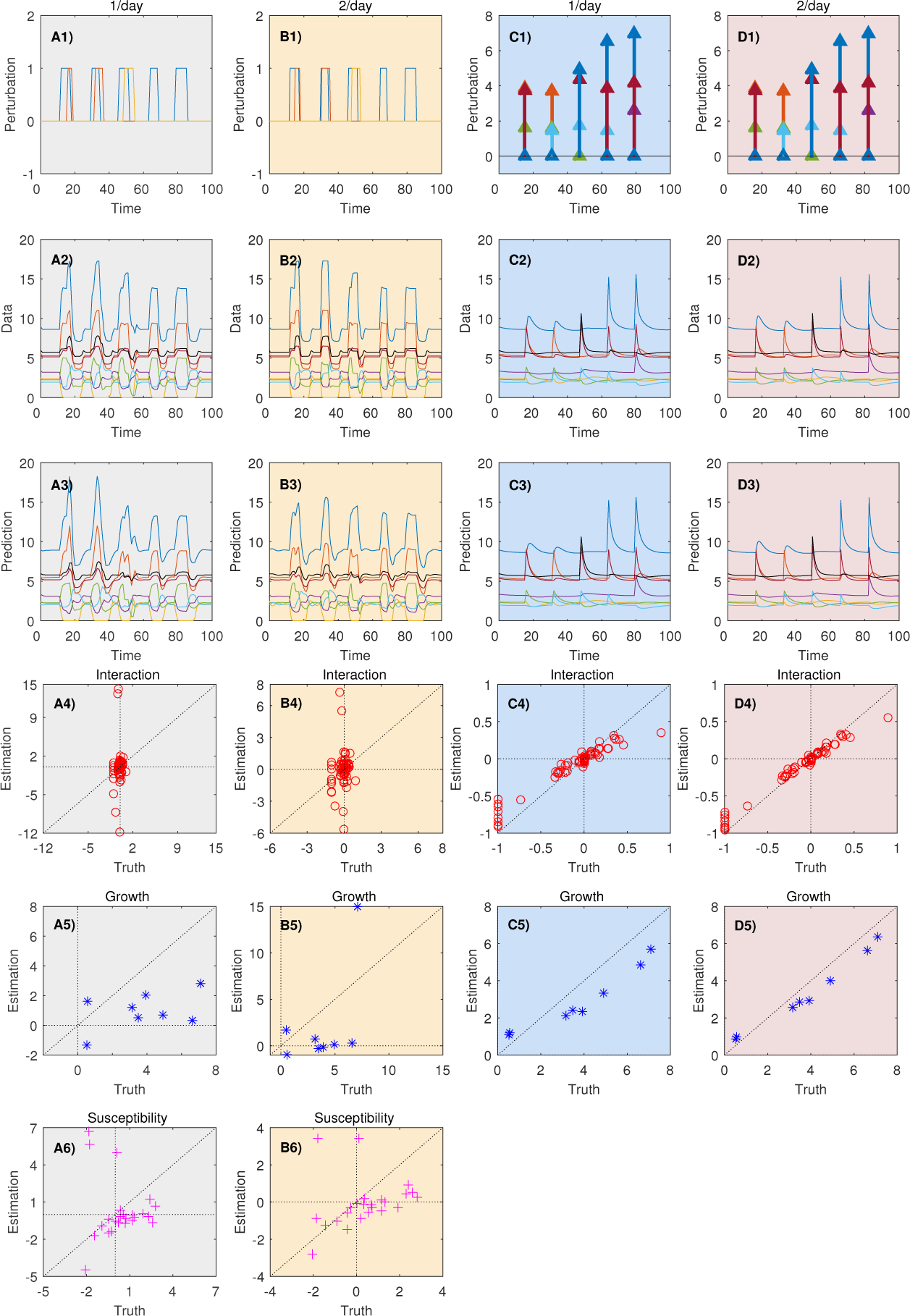
Inferred results of the same microbial system with two different driving response schemes under the same sampling rate. All the data are sampled from the same microbial systems. Column A represent the data and inferred results under binary perturbations sampled once per day. Column B is same as A but the sampling rate is doubled. Column C represent the data sampled once per day and the inferred results by changing the initial conditions only. Column D is the same as C but the sampling rate is doubled. Clearly we get better inference results by changing the initial conditions and doubling the sampling rate.

